# Laser capture microdissection in combination with mass spectrometry: Approach to characterization of tissue-specific proteomes of *Eudiplozoon nipponicum* (Monogenea, Polyopisthocotylea)

**DOI:** 10.1101/2020.03.31.018366

**Authors:** Pavel Roudnický, David Potěšil, Zbyněk Zdráhal, Milan Gelnar, Martin Kašný

## Abstract

*Eudiplozoon nipponicum* (Goto, 1891) is a hematophagous monogenean ectoparasite which inhabits the gills of the common carp (*Cyprinus carpio*). Heavy infestation can lead to anemia and in conjunction with secondary bacterial infections cause poor health and eventual death of the host.

This study is based on an innovative approach to protein localization which has never been used in parasitology before. Using laser capture microdissection, we dissected particular areas of the parasite body without contaminating the samples by surrounding tissue, and in combination with analysis by mass spectrometry obtained tissue-specific proteomes of tegument, intestine, and parenchyma of our model organism, *E. nipponicum*. We successfully verified the presence of certain functional proteins (e.g. cathepsin L) in tissues where their presence was expected (intestine) and confirmed that there were no traces of these proteins in other tissues (tegument and parenchyma). Additionally, we identified a total of 2,059 proteins, including 72 peptidases and 33 peptidase inhibitors. As expected, the greatest variety was found in the intestine and the lowest variety in the parenchyma.

Our results are significant on two levels. Firstly, we demonstrated how one can localize all proteins in one analysis and without using laboratory animals (antibodies for immunolocalization of single proteins). Secondly, this study offers the first complex proteomic data on not only the *E. nipponicum* but within the whole class of Monogenea, which was from this point of view until recently neglected.

## Introduction

Laser-capture microdissection (LCM) was developed as a powerful and reliable tool to overcome the heterogeneity of the specimen [1]. It enables isolation of specific tissues, individual cells, or even individual organelles from complex samples based on cell morphology [2,3]. The method was first described in the second half of the 20^th^ century and since then further developed and modified [2,4–7]. Nowadays, LCM combines laser excision with high-resolution microscopic control, which makes it possible to precisely capture for instance individual cells or even only nucleus while keeping track of the location and morphology of the source tissue [2,3,8].

LCM technology has been used in a wide variety of applications with focus on genomic, transcriptomic, and even proteomic analyses, such as dissection of polar bodies from oocytes for pre-fertilization genetic diagnosis [9], transcriptome-wide analysis of blood vessels from human skin and wound-edge tissue [10], proteomic profiling of dentoalveolar tissues [11], and many other areas [12–14]. But only one study so far used a combination of LCM and mass spectrometry (MS) to localize unique proteins, potential biomarkers, when dealing with the heterogeneity of breast tumor [15].

LCM has also been used in parasitology, especially in sample preparation. Recent studies employed this method in genome sequencing of *Plasmodium relictum*, where it was used to dissect the parasite from infected erythrocytes while excluding the host nucleus [16]. It has also been recently applied in a transcriptomic analysis of *Plasmodium cynomolgi* for dissection of hypnozoites from hepatocytes [17], in molecular analysis of *Henneguya adiposa* from the fins of catfish [18], and in investigation of ferritin gene expression in vitelline cells and tissue-specific gene profiling (gastrodermis, vitelline, and ovary tissue) of *Schistosoma japonicum* [19,20] and *Schistosoma mansoni* [21,22]. In a study focused on changes in protein composition in intermediate snail host, infected or uninfected by *S. mansoni*, the method was used to dissect hemocytes and sporocysts [23]. Up to now, however, this technique was not applied to protein localization, although analysis of tissue-specific proteomes could be an elegant way of localizing the molecules of interest, thus facilitating a better understanding of their function. It would be faster and less laborious to obtain results this way than by traditional approaches, such as immunolocalization or *in situ* hybridization, which require preparation of recombinant proteins, subsequent immunization processes, and development of RNA-probes.

With respect to investigation of the molecular content of the tissue within its morphological context, it was the technique of mass spectrometry imaging (MSI) that made it possible [24]. In parasitology, one study examined the chemical markers of the surface of *S. mansoni* by MSI to distinguish between the sexes and the strains [25], while another study dealt with the same organism and investigated the composition of internal organs by histological sections [26]. Because of the technical limits of MSI, in both cases, the focus was on the relatively small molecules of triacylglycerols and phosphatidylcholines, not on proteins. This demonstrates why we opted for a different approach: MSI is well-suited to the investigation of small molecules but lacks the ability to identify proteins whose size exceeds app. 15 or 25 kDa [27,28], depending on the specific instrumental setup.

The ability to capture higher molecular weights is essential in search for functional proteins, because their weight usually ranges around several tens of kilodaltons. For example, the digestive peptidases of *E. nipponicum*, such as like cathepsins L and B, have molecular weight of ~25 kDa, ~29 kDa, respectively [29], while one of the serine peptidase inhibitors from the same organism has molecular weight of ~45 kDa [30].

In the present study, our aim is first of all to investigate the potential of using laser capture microdissection (LCM) and liquid chromatography coupled to tandem mass spectrometry (LC-MS/MS) for protein localization and secondly, to obtain tissue-specific proteomes of our experimental organism *Eudiplozoon nipponicum* Goto, 1891 (Polyopisthocotylea). This monogenean is a common hematophagous ectoparasite which inhabits the gills of the common carp (*Cyprinus carpio*). For this species, as for the whole group of parasitic Monogenea, proteomic but also genomic and transcriptomic data are still scarce. In particular, no complex proteomic or secretomic data have so far been published at all and transcriptomic data are available only for *Neobenedenia melleni* (available in the NCBI BioSample database, http://www.ncbi.nlm.nih.gov/biosample/, accession number SAMN00169373). With respect to genomic data, the situation is somewhat better but only to two complete genomes are available, namely those of *Gyrodactylus salaris* [31] and *Protopolystoma xenopodis* (available in the NCBI BioProject database, https://www.ncbi.nlm.nih.gov/bioproject/, accession number PRJEB1201). For some monogenean species, mitochondrial genomes have, however, been mapped: *N. melleni* [32], *G. salaris* [33], *Gyrodactylus thymalli* [34], *Pseudochauhanea macrorchis* [35], *Benedenia hoshinai* [36], *Benedenia humboldti* [37], *Paratetraonchoides inermis* [38], *Lamellodiscus spari* and *Lepidotrema longipenis* [39], and *Thaparocleidus asoti* and *T. varicus* [40].

In recent years, this situation started to improve because *E. nipponicum* has been studied in a broader context. Several functional protein molecules of *E. nipponicum* were described [29,30,41–44] and the genome, transcriptome, and secretome of this organism are soon to be published. With this study, we significantly enrich available information on monogenean functional molecular biology by describing protein distribution in selected *E. nipponicum* tissues.

## Materials and Methods

### Parasite material: Collection and fixation

Adults of *E. nipponicum* were collected from freshly sacrificed specimens of *Cyprinus carpio* provided by Rybářství Třeboň a.s., Rybářská 801, Třeboň 379 01, Czech Republic. Isolation and taxonomic identification of the individual worms from the gills was performed as described previously [30].

Extracted worms were thoroughly washed in 10mM PBS pH 7.2 (PBS) to remove gill tissue debris. Then they were placed in a Petri dish and glass cover placed on them to keep them in stretched flat position. Solution of 4% paraformaldehyde in PBS was pipetted into the Petri dish and the sample was left in room temperature for 4hrs. After fixation, samples were rinsed with PBS buffer and transferred into cryofixation molds, which were then immersed in OCT compound (Tissue-Tek^®^) and left for 1 hr in room temperature. The molds were then placed in dry ice and frozen blocks stored at −80°C ([45], modified).

### Ethics statement

All procedures performed in studies involving animals were carried out in accordance with European Directive 2010/63/EU and Czech laws 246/1992 and 359/2012 which regulate research involving animals. All experiments were performed with the legal consent of the Animal Care and Use Committee of Masaryk University and of the Research and Development Section of the Ministry of Education, Youth, and Sports of the Czech Republic.

### Cryosectioning

The OCT-embedded worms were sectioned in a cryotome (Leica CM1900 UV, −20°C) in 12 μm thick slices, which were placed on microdissection slides (MembraneSlide 1.0 PEN, Carl Zeiss). Prior the LCM, OCT surrounding the tissues had to be removed, which was achieved by careful dipping of the slides in ddH2O. In the next step, the tissues were dehydrated by EtOH (96%, two washes for 30 s) and the slides either subjected to LCM immediately or stored in 4°C for a short time. During cryosectioning, we placed ten to twelve worm sections on a single membrane slide.

### Laser capture microdissection

Target tissues were extracted from the histological sections using microdissector PALM MicroBeam (Carl Zeiss) controlled by software PALMRobo 4.6 Pro (Carl Zeiss). Dissected samples were collected into special caps filled with adhesive material for dry collection (AdhesiveCap 500, Carl Zeiss). Cutting options (laser intensity and focus) were adjusted for each type of tissue specifically due to the nature of the parasite body. Only limited, well distinguished areas were dissected at once for individual tissue types (concretely parenchyma up to 2,000 μm^2^ per cut, intestine up to 350 μm^2^ per cut, tegument up to 500 μm^2^ per cut) with overall area of 1 mm^2^ per tissue type. We prepared and analyzed two full sets of tissue samples (biological replicates).

### Sample preparation for LC-MS/MS

Microdissected tissue samples were lysed in SDT buffer (4% SDS, 0.1M DTT, 0.1M Tris/HCl, pH 7.6) in a thermomixer (Eppendorf ThermoMixer^®^ C, 30 min, 95°C, 750 rpm). After that, samples were centrifuged (15 min, 20,000 x g) and the supernatant used for filter-aided sample preparation as described elsewhere [46] using 0.5 μg/sample of trypsin (sequencing grade, Promega). Resulting peptides were used for LC-MS/MS analyses. Total peptide amount after the sample preparation was estimated using LC-MS analysis on RSLCnano system online coupled with HCTUltra ion trap (Bruker Daltonics) based on the area under the total ion current curve, using MEC cell line tryptic digest as external calibrant.

### LC-MS/MS analysis and data evaluation

LC-MS/MS analyses of all peptide mixtures were done using UltiMate™ 3000 RSLCnano system connected to Orbitrap Fusion Lumos Tribrid spectrometer (Thermo Fisher Scientific). Prior to LC separation, tryptic digests were online concentrated and desalted using trapping column (X-Bridge BEH 130 C18, dimensions 30 mm × 100 μm, 3.5 μm particles; Waters). After washing of trapping column with 0.1% FA, the peptides were eluted in backflush mode (flow 0.3 μl/min) from the trapping column onto an analytical column (Acclaim Pepmap100 C18, 3 μm particles, 75 μm × 500 mm; Thermo Fisher Scientific) during 130 min gradient (1—80 % of mobile phase B; mobile phase A: 0.1% FA in water; mobile phase B: 0.1% FA in 80% ACN).

MS data were acquired in a data-dependent mode, selecting up to 20 precursors based on precursor abundance in the survey scan. The resolution of the survey scan was 120,000 (350— 2000 m/z) with a target value of 4×10^5^ ions and maximum injection time of 100 ms. MS/MS spectra were acquired with a target value of 5×10^4^ ions (resolution 15,000 at 110 m/z) and maximum injection time of 22 ms. The isolation window for fragmentation was set to 1.2 m/z.

For data evaluation, we used MaxQuant software (v1.6.2.10) [47] with inbuild Andromeda search engine [48]. Searches against in-house made protein databases were undertaken: *E. nipponicum* (37,076 sequences, manuscript in preparation), *C. carpio* (63,928 sequences, based on https://www.ncbi.nlm.nih.gov/genome/?term=10839), and cRAP contaminants (based on http://www.thegpm.org/crap). Modifications for all database searches were set as follows: oxidation (M), deamidation (N, Q), and acetylation (Protein N-term) as variable modifications, with carbamidomethylation (C) as a fixed modification. Enzyme specificity was tryptic with two permissible miscleavages. Only peptides and proteins with false discovery rate threshold under 0.01 and proteins with at least two peptide identification were considered. Relative protein abundance was assessed using protein intensities calculated by MaxQuant. For the purpose of this article, protein groups reported by MaxQuant are referred to as proteins. A complete list of proteins within each protein group can be viewed in the supporting material (S1, Table). Mass spectrometry proteomics data were deposited to the ProteomeXchange Consortium via PRIDE [49] partner repository under dataset identifier PXD017275.

Intensities of reported proteins were further evaluated using software container environment (https://github.com/OmicsWorkflows/KNIME_docker_vnc; version 3.7.1a). Processing workflow is available upon request: it covers decoy hits and removal of contaminant protein groups (cRAP), protein group intensities log2 transformation and normalization. Carp proteins were filtered out as contaminants. Mapping to *E. nipponicum* transcriptome (accession GFYM00000000.1) was also undertaken, which enriched protein identifications by annotations (GO terms, MEROPS, Pfam and InterPro accessions).

### Comparative analyses

Datasets obtained from the three *E. nipponicum* tissues (intestine, parenchyma, and tegument) were compared with each other to identify common and unique proteins, with special focus on inhibitors and peptidases. Annotations of proteins unique to each tissue were loaded from the Kyoto Encyclopedia of Genes and Genomes (KEGG) while excluding the “organismal system” and “human disease” categories [50]. Based on the measured intensity of proteins found in each tissue, the most abundant peptidases and inhibitors were identified using fold change ratios calculated between the respective tissue samples.

### Localization of functional proteins

To assess whether our approach could also be applied to protein localization, we compared proteins identified in each tissue with previously characterized proteins whose functions were described in recent publications: cathepsins L and B [29], stefin [43], kunitz [42] and serpin [30].

## Results

### Laser capture microdissection of tissue samples

We targeted 3 types of tissue — intestine, tegument and parenchyma (Fig 1). These tissues samples were cut out from 10 μm thick longitudinal cryosections of several individuals of *E. nipponicum*. Total area of each tissue sample was 1 mm^2^, which corresponds to volume of app. 0.012μm^3^.

**Fig 1.**
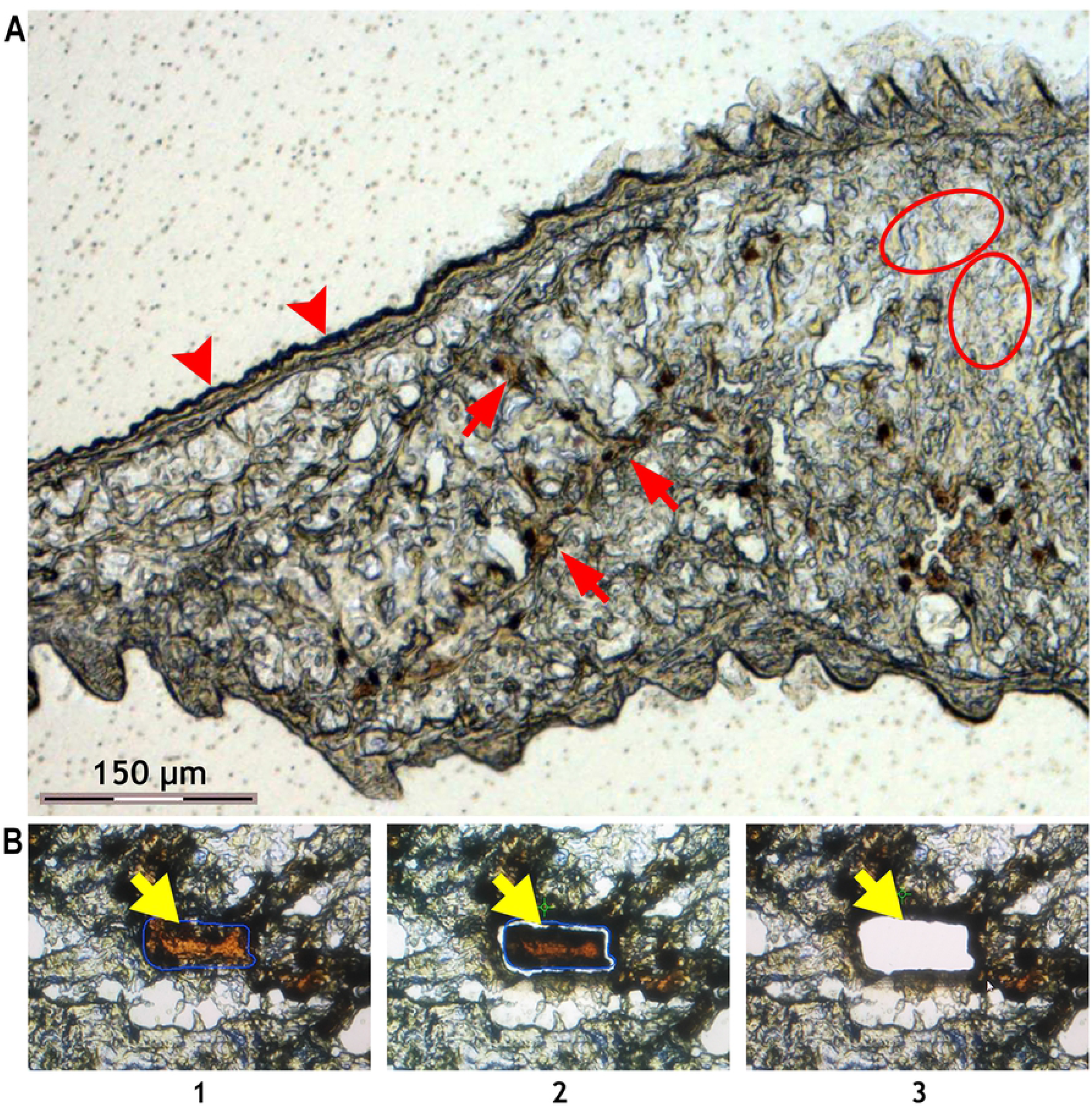
Laser microdissection of chosen tissues of E. nipponicum. (A) Part of a cryosectioned body. Branched intestine filled with host blood is clearly visible (red arrow). Tegument is visible at the edge of the section (red arrowhead). Parenchyma is the area with no visible organelles (red ovals). (B) Process of microdissection: 1) selection of desired area; 2) laser section; 3) catapulting specimen to collection tube. Yellow arrows point to the dissected area.

### LC-MS/MS analyses and data evaluation

The total peptide yield estimation after the filter aided sample preparation (FASP) sample processing step showed there were app. 40 μg in intestine, 8 μg in tegument and 6 μg in parenchyma in each dissected tissue type of comparable size, suggesting higher overall protein content in intestine. In total, i.e. in all three tissues jointly, we identified 2,059 proteins. Of these proteins, 1,978 were found in the intestine, 1,425 in the tegument, and 1,302 in the parenchyma. An overview is listed in Table 1, while Fig 2 shows how many proteins are unique or shared among the tissues. There were also 273 carp proteins identified, which were filtered out and they are not included in results.

**Table 1.**
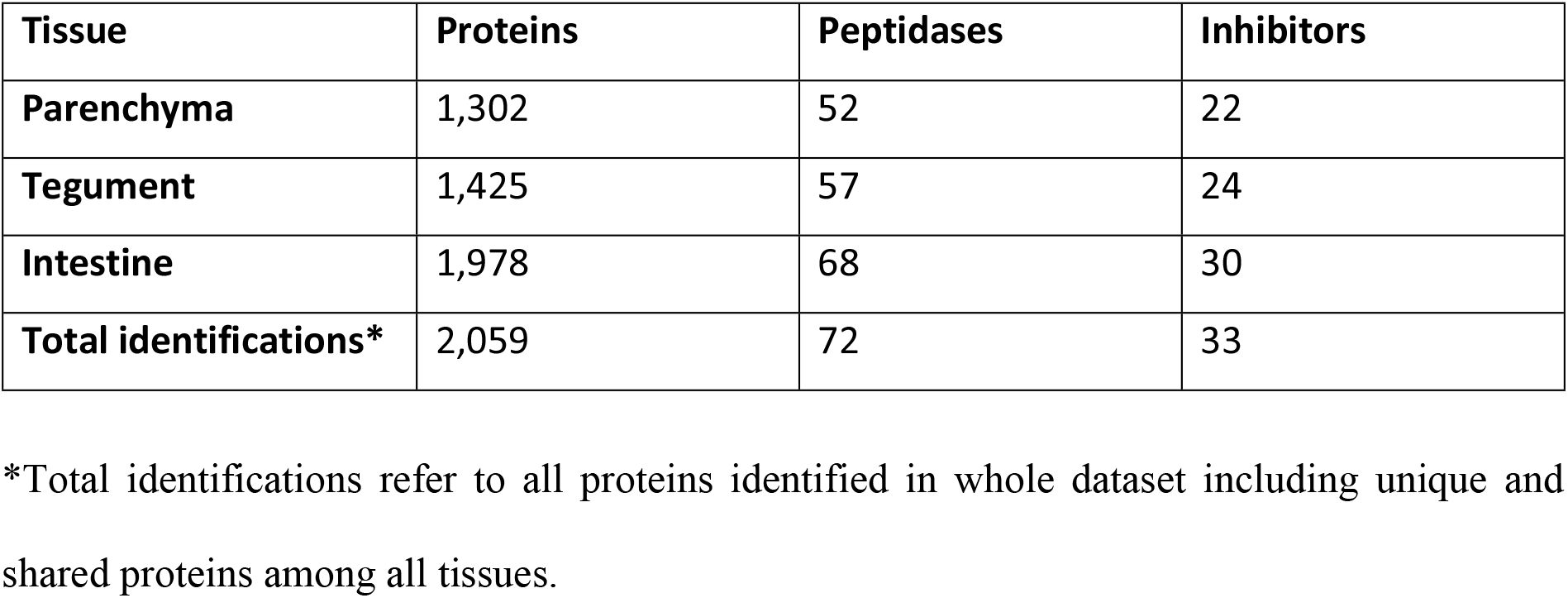
Resulting numbers of identified proteins in each tissue.

**Fig 2.**
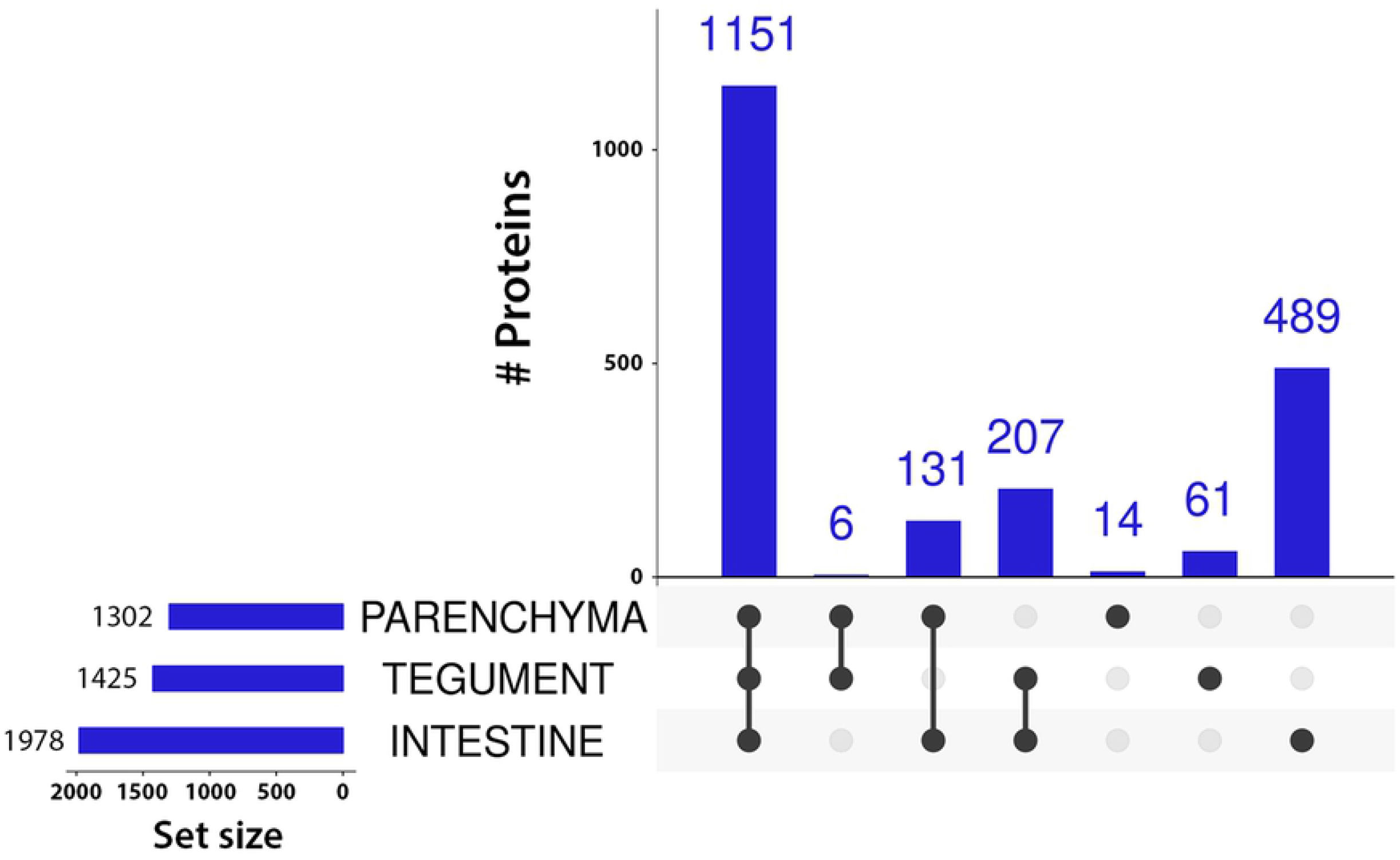
UpSet plot showing numbers of identifications in each tissue. Only proteins identified by ≥ 2 peptides in at least one replicate are considered to be present. Intersections show proteins present in all tissues marked by connected black dots. Black dots without connections refer to proteins unique to that tissue.

Unique proteins from each tissue were assigned to KEGG pathways (Fig 3). Most annotations turned out to be in the intestine dataset (250 out of 489 entries; 51.1%), followed by the tegument (20 out of 61; 32.8%) and the parenchyma (4 out of 14; 28.6%).

**Fig 3.**
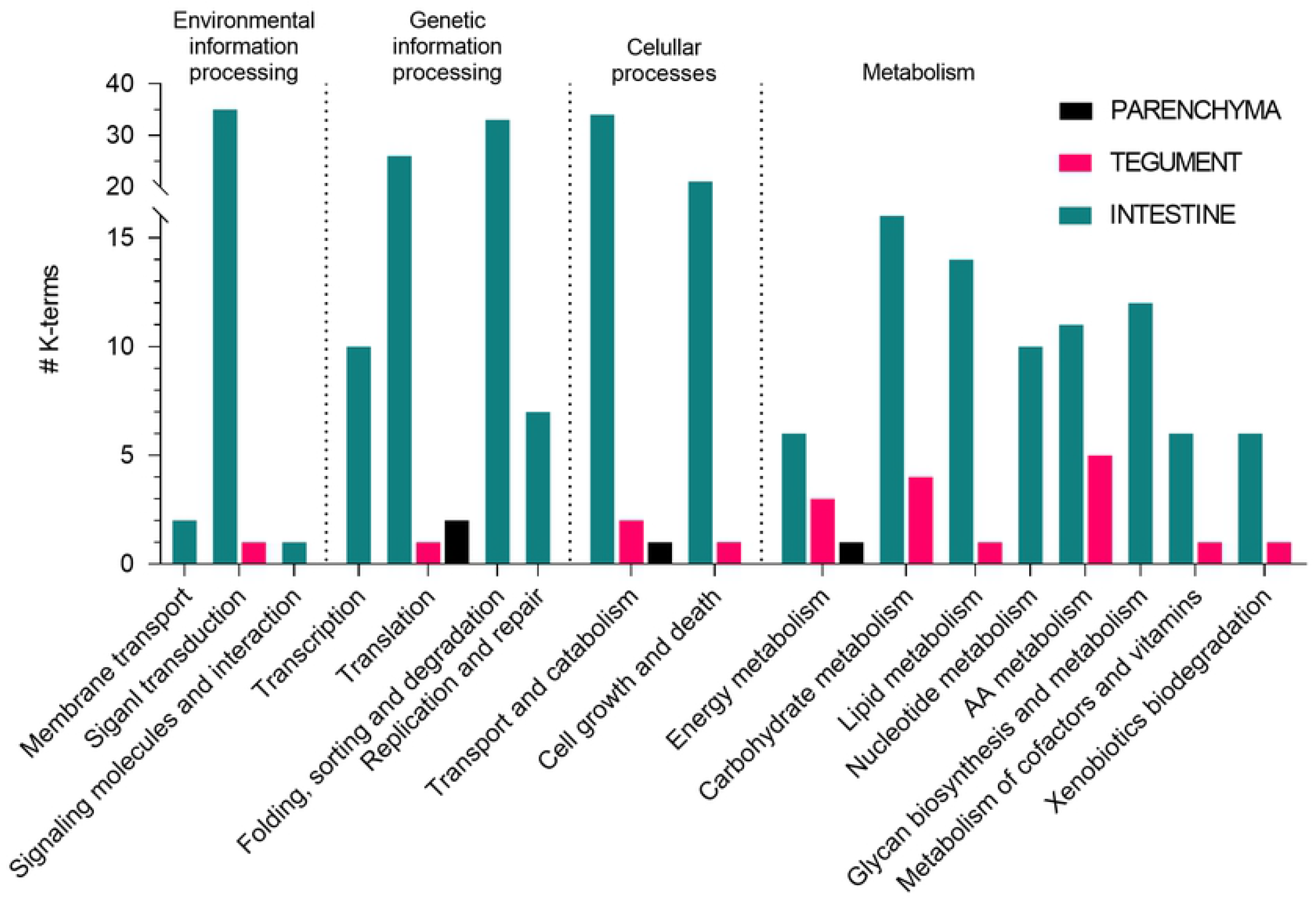
KEGG pathway annotation of the unique proteins from each tissue. Only proteins identified by ≥ 2 peptides in both replicates of a given sample and by ≤ 1 peptide in other samples are considered unique to a sample.

With respect to **peptidases**, we were able to identify 72 in all three tissues jointly. Numbers of peptidases identified in each tissue are shown in the Fig. 4. Most peptidases were found in the intestine, where one also finds most of the unique ones.

**Fig 4.**
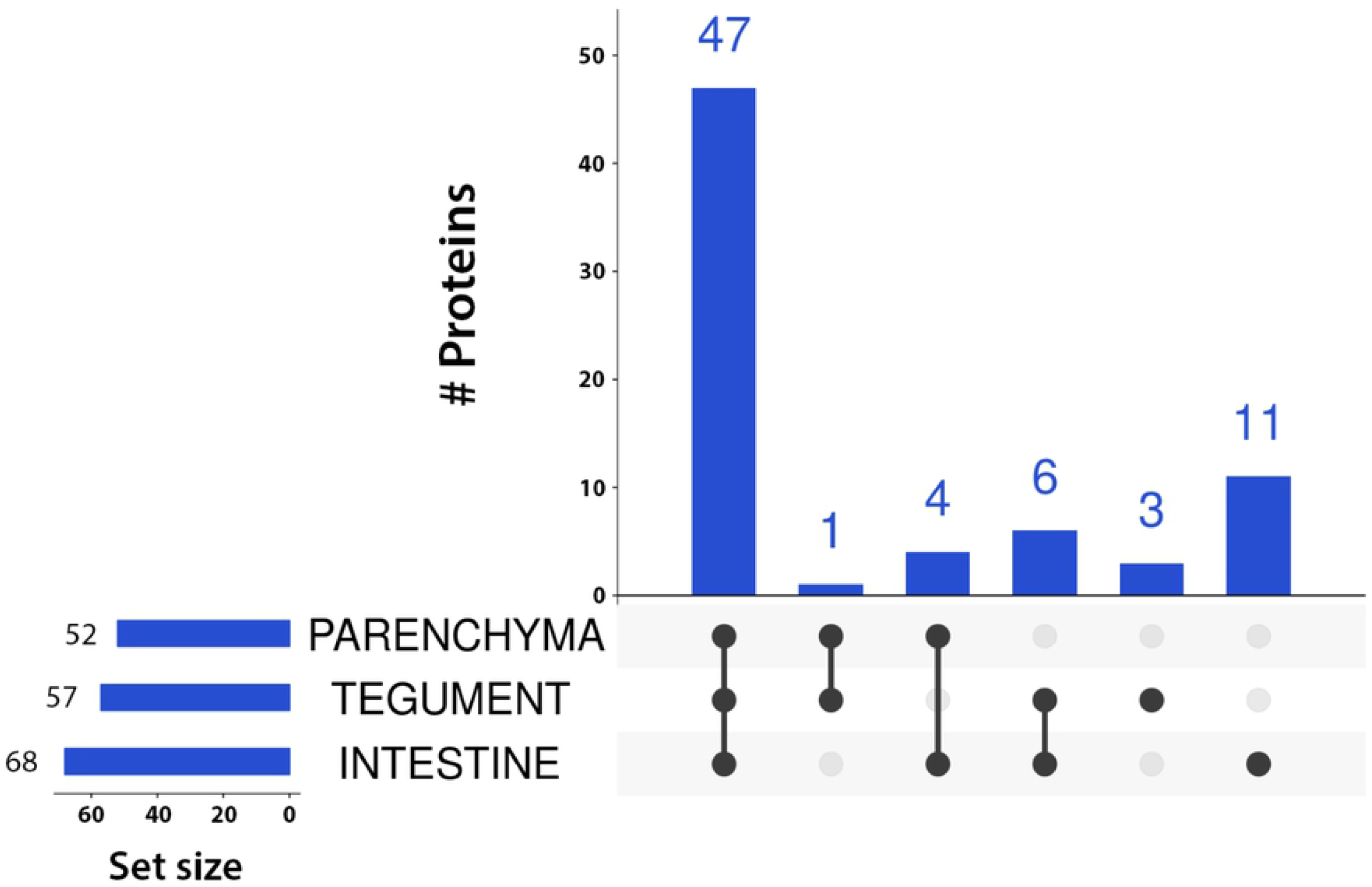
UpSet plot showing the number of peptidases in each tissue. Only proteins identified by ≥ 2 peptides in at least one replicate are considered to be present. Proteins common to several of tissues are marked by connected black dots. Black dots without connections refer to proteins unique to that tissue.

Peptidases unique to a tissue were classified into several groups based on their catalytic mechanism. There are four catalytic groups present in our dataset and only peptidases from the intestine belong to all of them, i.e. to serine, aspartic, cysteine, and metallo. Tegumental peptidases belong to two catalytic groups (serine and metallo), while in the parenchyma, we found no unique peptidases (Fig 5).

**Fig 5.**
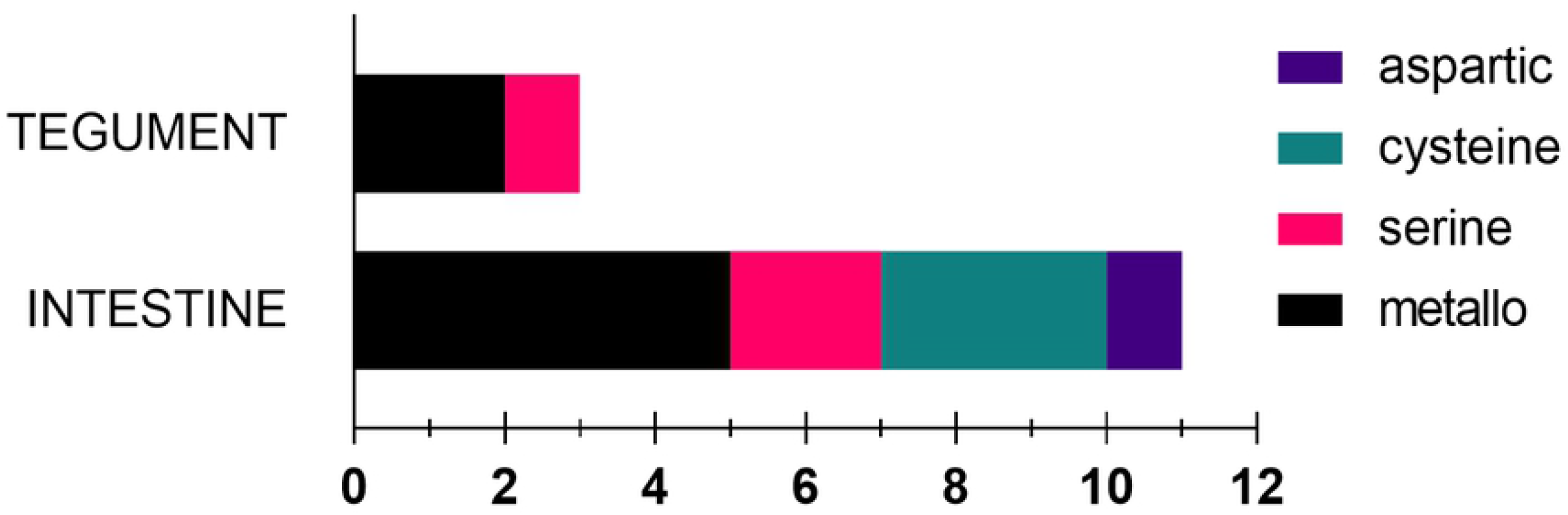
Catalytic groups of unique peptidases. Only proteins identified by ≥ 2 peptides in both replicates of a sample and by ≤ 1 peptide in other samples are considered unique to a tissue.

Furthermore, we identified 33 **inhibitors** in all three tissues jointly. Numbers of inhibitors identified in each tissue are shown in Fig 6. Like in the peptidases, most inhibitors are present in the intestine.

**Fig 6.**
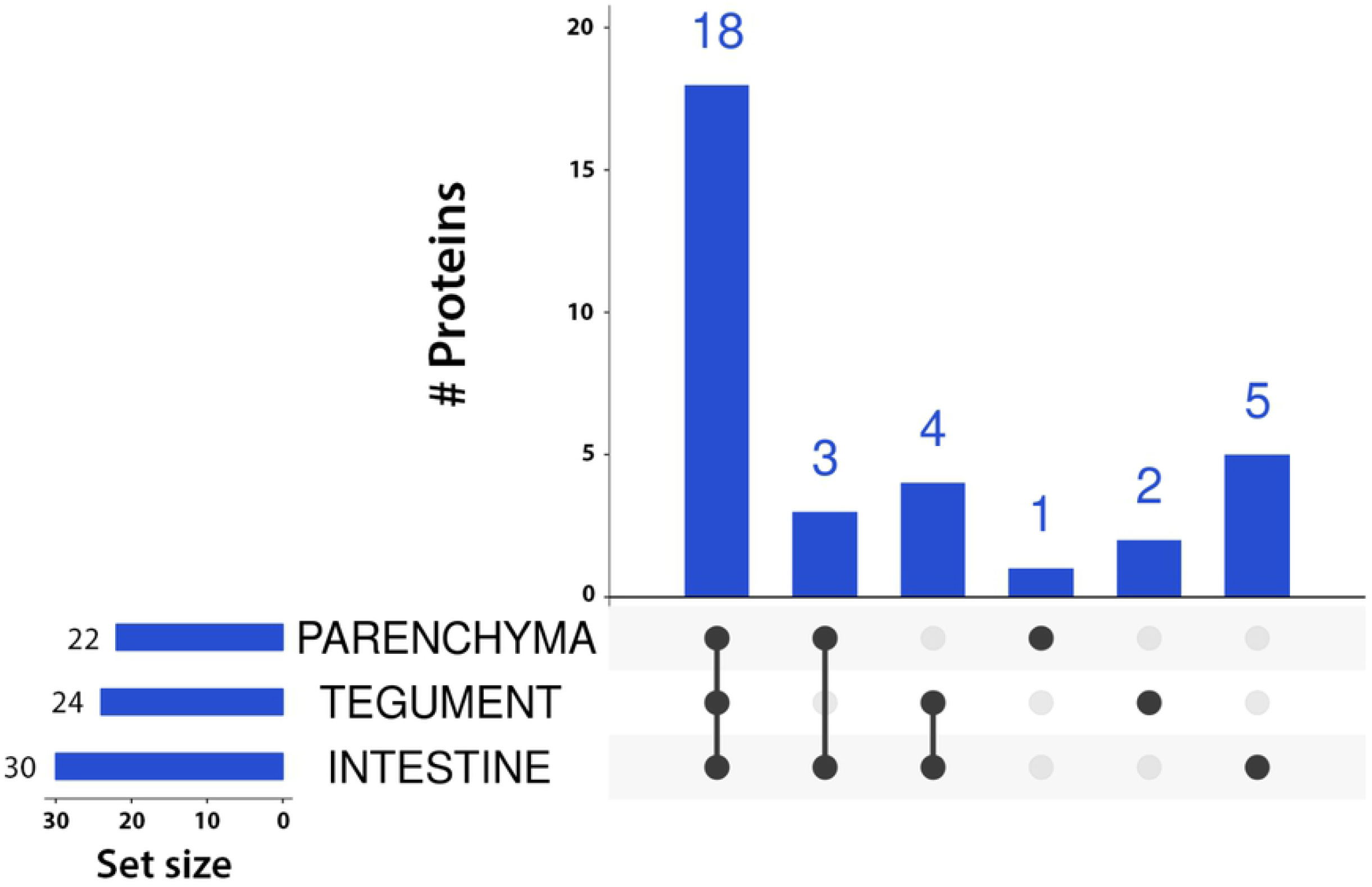
UpSet plot showing the number of protein inhibitors found in each tissue. Only proteins identified by ≥ 2 peptides in at least one replicate are considered to be present.

Intersections show proteins common to tissues connected by black dots. Black dots without connections refer to proteins unique to that tissue.

Unique inhibitors were sorted in several families according to MEROPS (Fig 7). Inhibitors from the intestine were again the most varied ones, covering four (I2, I25, I29, and I32) of the seven families present. In the parenchyma, only one family was present (I4), while tegumental inhibitors belonged to two families (I21 and I87).

**Fig 7.**
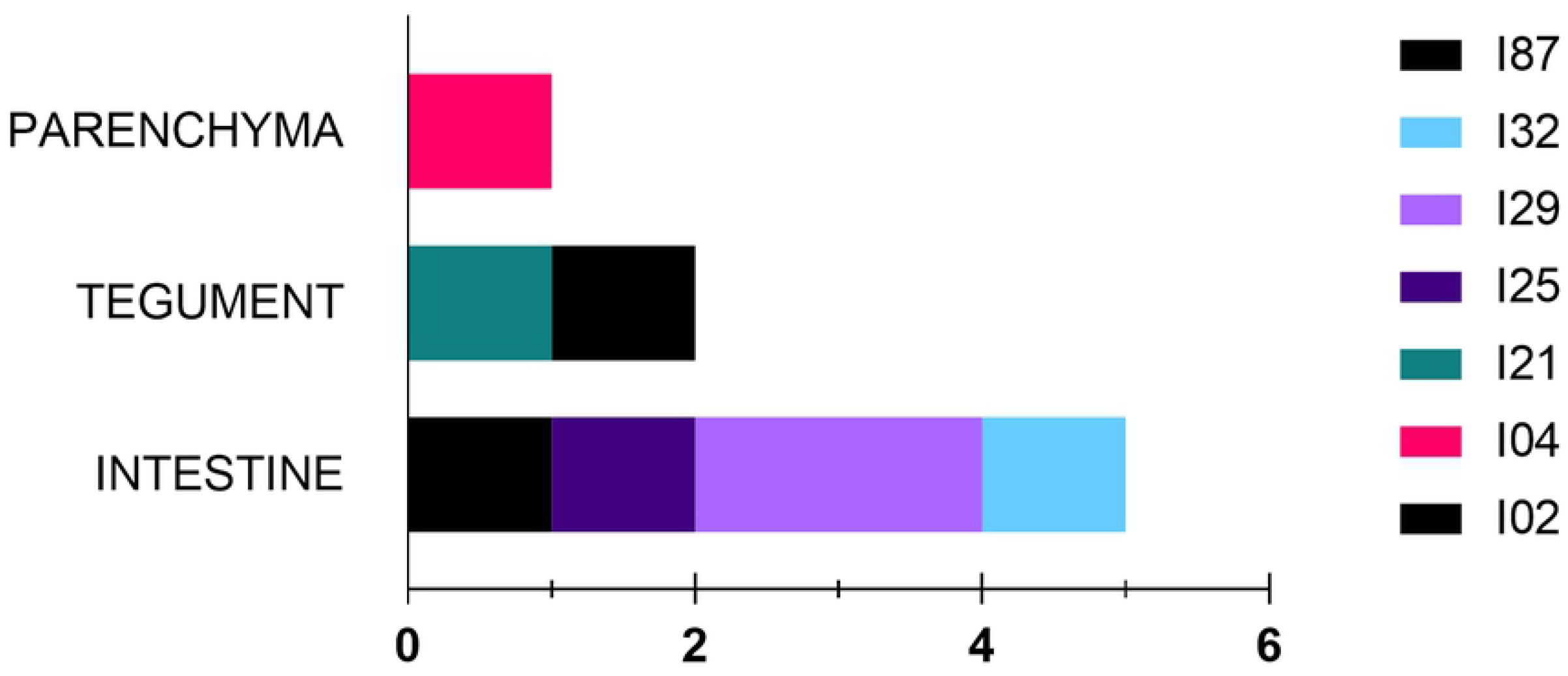
A distribution of unique protein inhibitors into families (MEROPS). Only proteins identified by ≥ 2 peptides in both replicates of a sample and by ≤ 1 peptide in other samples are considered unique to a tissue.

### The differently expressed peptidases and inhibitors

Subsequently, we compared the abundance of inhibitors and peptidases in each tissue based on protein intensities. We found three peptidases more abundant in the intestine and four in the tegument. With respect to inhibitors, we found one more abundant in the intestine and one in the parenchyma. For detailed information, see Table 2.

**Table 2.**
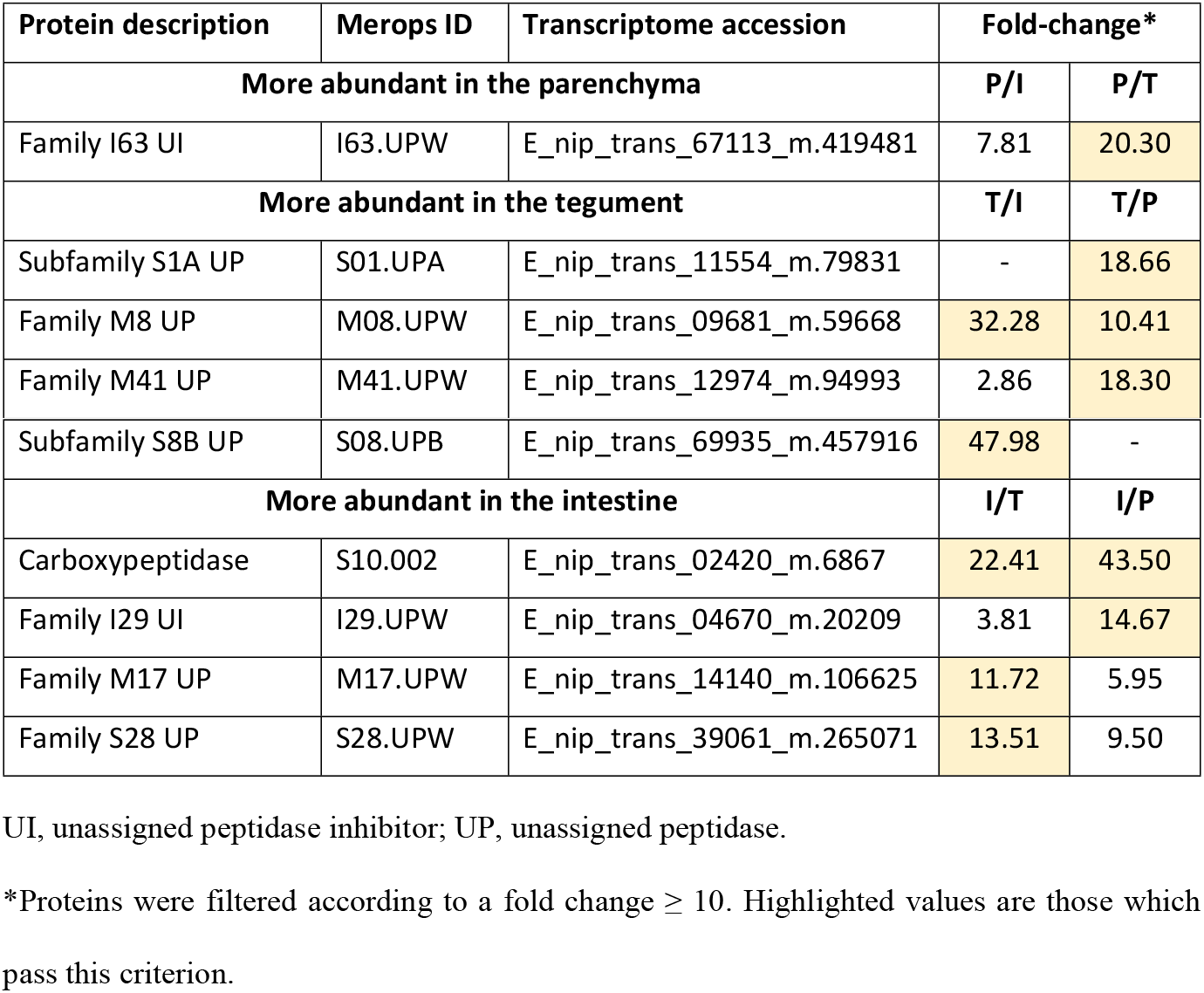
List of the most abundant peptidases and inhibitors.

| Protein description | Merops ID | Transcriptome accession | Fold-change* |  |
| --- | --- | --- | --- | --- |
| More abundant in the parenchyma |  |  | P/I | P/T |
| Family I63 UI | I63.UPW | E_nip_trans_67113_m.419481 | 7.81 | 20.30 |
| More abundant in the tegument |  |  | T/I | T/P |
| Subfamily S1A UP | S01.UPA | E_nip_trans_11554_m.79831 | - | 18.66 |
| Family M8 UP | M08.UPW | E_nip_trans_09681_m.59668 | 32.28 | 10.41 |
| Family M41 UP | M41.UPW | E_nip_trans_12974_m.94993 | 2.86 | 18.30 |
| Subfamily S8B UP | S08.UPB | E_nip_trans_69935_m.457916 | 47.98 | - |
| **More abundant in the intestine** |  |  | **I/T** | **I/P** |
| Carboxypeptidase | S10.002 | E_nip_trans_02420_m.6867 | 22.41 | 43.50 |
| Family I29 UI | I29.UPW | E_nip_trans_04670_m.20209 | 3.81 | 14.67 |
| Family M17 UP | M17.UPW | E_nip_trans_14140_m.106625 | 11.72 | 5.95 |
| Family S28 UP | S28.UPW | E_nip_trans_39061_m.265071 | 13.51 | 9.50 |

UI, unassigned peptidase inhibitor; UP, unassigned peptidase.
\*Proteins were filtered according to a fold change $\geq 10$ . Highlighted values are those which pass this criterion.

### Localization of functional proteins

Seven cathepsins (CL3, CL4, CL6b, c, d, e, and CB) were identified only in the intestine, two (CL1, CL5) in all tissues, and two have not been identified at all although based on a related publication [29], they should be present in the intestine as well. Both inhibitors, i.e. EnStef and EnSerp1, were present in all tissues (see Table 3).

**Table 3.**
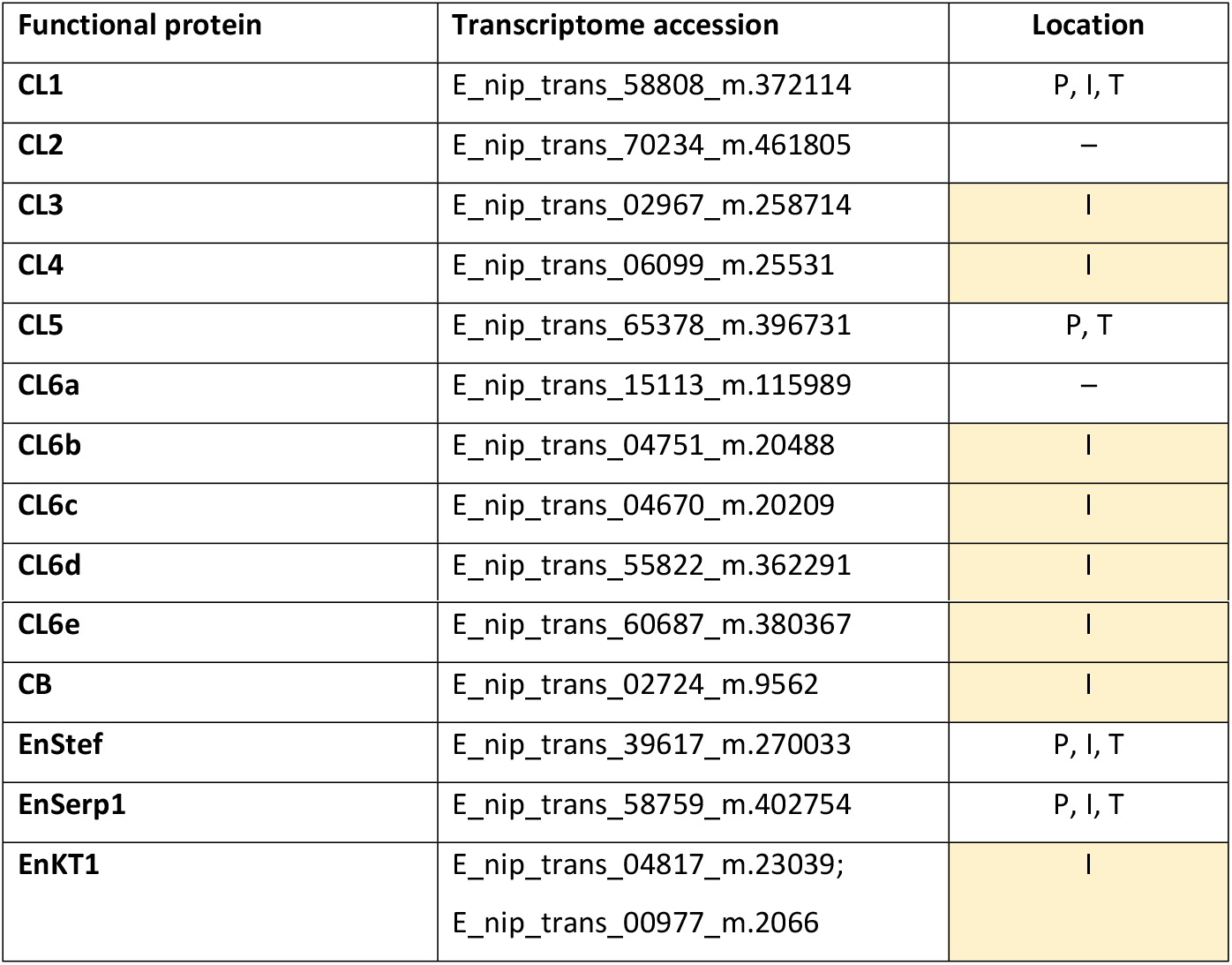
Localization of functional proteins.

## Discussion

In this study, we report the first tissue-specific proteomic analysis of parasitic monogenean *Eudiplozoon nipponicum*. Using laser capture microdissection, we dissected particular areas of three different tissues of the parasite body and the dissected samples were subjected to mass spectrometry-based proteomic analysis. This allowed us to characterize tissue-specific proteomes and to assess the suitability of this approach for proteins tissue-localization. We chose to focus on intestine, tegument and parenchyma because those tissues can easily be distinguished without any additional histological staining. Moreover, functional proteins from the intestine are of great importance due to their involvement in host-parasite interplay and a better understanding of these processes could help contribute to a control of infection not only by *E. nipponicum* but possibly other monogeneans as well. However, only a handful of studies so far focused on particular functional proteins [29,30,42–44] and the more complex proteomic data have been missing until now.

In this study, we identified 2,059 proteins. Most identifications were made in the intestine sample (1,978), less in the tegument (1,425), and the lowest number of identifications was made in the parenchyma sample (1,302) (Table 1, Fig 2). Of these proteins, 72 (3.50%) were peptidases and 33 (1.60%) inhibitors (Table 1), which is in good agreement with ratios found in other parasites, such as the fluke *S. mansoni* proteome on UniProt (UP000008854, [51]), which contains 6,858 entries and the MEROPS lists 284 (4.14%) peptidases and 129 (1.88%) inhibitors, or the tick *Ixodes ricinus* proteome (UP000001555, Caler et al. 2008 not published), which contains 20,473 entries and the MEROPS lists 318 (1,55%) peptidases and 277 (1.35%) inhibitors (MEROPS database accessed January 18, 2020). Apart from this, we identified also 273 carp proteins. The vast majority were part of the intestine sample set and related to food content as attested by the fact that we found blood proteins (hemoglobin subunits, heme-binding protein, complement factors, antihemorrhagic factor etc.). These proteins were filtered out as expected contaminants.

In terms of tissue specificity of peptidases and inhibitors (Fig 4 and 6), the richest environment is the intestine, which is the most metabolically active: both digestion and nutrient uptake take place in the gut lumen and hematin cells of the gastrodermis [29,52]. The tegument ended up the second, which could be due to the fact that this tissue protects the rest of the body from the environment. It is the site of various important processes, such as glycocalyx maintenance [53,54], sensory perception [55], and secretion [56]. The lowest variety of proteases and inhibitors was found in the parenchyma, where no major organs were dissected. This tissue served primarily as a control for the other two. KEGG pathway annotations provided further support to these conclusions (Fig 3), although the input (unique proteins) for analysis was different for each tissue. Nonetheless, with respect to for instance metabolism, we can conclude that the tegument is much more frequently represented in the particular sections (such as carbohydrate and amino acid metabolism), which may be related to its abovementioned function in surface maintenance. On the other hand, the lack of uniqueness in the parenchyma supports our initial expectations, because this tissue provides skeletal and structural support and functions as nutrient transport and storage [57].

The catalytic groups, which cover the unique identified peptidases, are shown in Fig 5. In total, we found representatives of four groups. Most represented were serine, cysteine, and metallopeptidases. This correlates with the number of unique inhibitors, which were assigned to seven families, including representatives of metallo (I87), cysteine (I29, I25, I4), and serine (I4 and I2) peptidase inhibitors (Fig 7). In all these statistics, the intestine was once again dominant with respect to variety. This is once again related to its function, namely feeding, since most peptidases and inhibitors play a role in digestion, inhibition of blood coagulation, and prevention of inflammation [29,42,52].

This is reflected in case of the most abundant peptidases/inhibitors (Table 2). For example, in the intestine we found serine carboxypeptidase. Serine carboxypeptidases from family S10 are known to have a lysosomal function [58], but it is also assumed that they have a role in blood degradation. The latter function seems to be supported by findings on *S. japonicum*, where it has an expression profile similar to the digestive cathepsin [59], while carboxypeptidase, its ortholog from tick *Ixodes ricinus*, has been described as a digestive enzyme [60]. There is also one abundant cysteine peptidase inhibitor from family I29. Its ortholog, a propeptide of *Fasciola hepatica* cathepsin L, proved to be a potent and selective inhibitor which plays a role in cathepsin L maturing and activity regulation [61], which makes its role similar to the *E. nipponicum* cysteine inhibitor.

In the tegument, we found the most abundant serine peptidase from the S8 family of diverse peptidases, most of which play a housekeeping role in protein turnover or processing of precursors of bioactive proteins [62].

In the parenchyma, one functional protein is significantly abundant: an inhibitor from the I63 family. This family contains an inhibitor of metallopeptidase pappalysin-1, which promotes cell growth [63]. We can therefore speculate that the identified inhibitor may play a role in the regulation of a peptidase with similar function.

Moreover, since we have access to the preliminary version of secretome of *E. nipponicum* (manuscript in preparation), we compared our data with it and saw that most secretome proteins are in fact a subset of data obtained in the present study. This is unsurprising because proteins in the secretome are produced by the parasite’s glands and intestine and at the time of production, they are situated in the parasite body, which is why they were detected by mass spectrometry in our analysis.

Finally, one of the main goals of this study was to investigate the potential of application of this methodology (LCM + LC-MS/MS) for protein localization. In this approach, one can select just those organs, tissues, or cells one is interested in and they can be extracted cleanly, that is, without contamination by unwanted surrounding tissue. And while this approach also requires further fine-tuning, it is markedly less laborious and time-consuming than the generally used techniques of immunolocalization and *in situ* hybridization [64]. Positive is also the fact that the procedure dispenses with the need to use laboratory animals. In the end, once the analysis is finished, one can go over the list of identified proteins and see where a protein of interest was identified. This can be done repeatedly and for different proteins without the need for additional experiments. The results of this study show that this method confirmed the localization of previously characterized cathepsins and kunitz of *E. nipponicum* [29,42] (Table 3). On the other hand, there is still room for improvement because two of the eleven examined cathepsins were not identified and another two were identified in tissues where they were not expected.

## Acknowledgements

We would like to thank Anna Pilátová for English proofreading.

## Supporting information

**S1 Table. LC-MS/MS analysis of *E. nipponicum* tissues**.

